# Intermediate heterogeneity modulates coupling between chain compaction and structure formation during protein folding

**DOI:** 10.1101/2025.11.26.690716

**Authors:** Anushka Kaushik, Jayant B. Udgaonkar

## Abstract

Polypeptide chains undergo both compaction and structure formation during folding, but the extent to which these processes are mechanistically coupled remains unclear. Although initial chain collapse can precede structure formation, the two processes invariably appear coupled at later stages of folding. This raises the question of whether the fraction of molecules that undergo initial collapse, as well as the degree of coupling between compaction and structure formation later during folding, are regulated by sequence-encoded structural constraints. To examine this, the folding of the small protein monellin was investigated using time-resolved fluorescence resonance energy transfer (trFRET) analyzed with the maximum entropy method to resolve sub-populations of molecules with native-like and unfolded-like dimensions. Mutation of Pro41 to Ala, or Pro93 to Ala, which relieve local backbone rigidity, selectively stabilized hidden minor conformations within the initial and later intermediate ensembles, respectively. In each case, the minor conformation had a segment that was more compact than in the major one, and its stabilization increased the number of molecules undergoing specific contraction to form the intermediate ensemble, without altering the extent of structure formation. Consequently, sub-populations within these intermediate ensembles could undergo chain contraction independently of structure formation. These findings identify intermediate-state heterogeneity, modifiable by backbone rigidity, as the basis for tunable coupling between chain compaction and structure formation during protein folding.

**Significance Statement:** Chain compaction and structure formation are central features of protein folding, yet how these two processes are coupled remains unclear. Although early chain collapse can precede structure formation, the two often appear coupled at later stages, raising the question of whether this coupling is obligatory or tunable. Here, the coupling is shown to be tunable. Using a quantitative framework that resolves coexisting compact intermediates, this study demonstrates that proline-imposed backbone rigidity governs intermediate-state heterogeneity, which in turn determines the separability of these processes.

## Introduction

The folding of a protein is a complex process (1–3), involving both compaction of the polypeptide chain (4–11) and the formation of secondary and tertiary structure (12, 13). The two processes are intimately linked, and seem to proceed together for the most part, but not much is known about how they are coupled to each other. Chain compaction favors the formation of secondary structure which is itself compact (5, 14, 15), and the coalescence of secondary structural units also induces compaction (16, 17). Different extents of chain compaction and structure formation occur during the early and late stages of the folding reaction and understanding how these changes are linked mechanistically to each other, remains an important challenge.

For many proteins, folding appears to begin with a collapse of the polypeptide chain, with (18–22) or without (23–26) the concurrent formation of structure. In the case of several proteins, different structural segments appear to undergo significantly different extents of initial collapse, as well as of later contraction (11, 27–30). Moreover, when initial collapse occurs, it does not happen to the same extent in all molecules (31–34). Hence, understanding how compaction is coupled to structure formation is complicated because the product of initial collapse is heterogeneous with sub-populations of molecules that have undergone substantial differential compaction across multiple structural segments, in dynamic equilibrium with sub-populations that have undergone only minor compaction (35). Delineating the heterogeneity inherent in the initial collapse reaction, as well as in the subsequent chain contraction that occurs as structure develops, is important for understanding how folding begins and proceeds.

Here, initial chain collapse and further chain contraction of single chain monellin (MNEI), a well-established model for both experimental (28, 29, 33–38) and computational (30, 39) studies of protein folding, have been studied. The folding of MNEI is known to begin with a non-specific chain collapse complete within 37 μs, which reduces the average diameter by approximately 30%, and leads to the formation of some non-native interactions (29). Subsequent stepwise consolidation of the collapsed globule occurs during the first millisecond of folding, without a further decrease in average dimensions, and results in the formation of a kinetic molten globule intermediate. At this stage, no significant secondary structure has formed, and even by 100 ms of folding in strongly native-like conditions, only about 10 % of N-like secondary structure is observed (28, 33).

Time-resolved fluorescence resonance energy transfer (trFRET) measurements, analyzed by the Maximum Entropy Method (MEM) (34, 35, 37, 40), not only are able to distinguish but can also quantify sub-populations of molecules present in different conformations. This methodology had been used earlier to monitor distances spanning different segments of the protein: the core (C), the β-sheet (B), the end-to-end distance (E), and the sole helix (H) of the protein (35). The H segment spanning the helix contracted gradually during folding. The distances spanning the C, B and E segments exhibited bimodal distributions at 100 milliseconds of folding, corresponding to two distinct sub-populations, unfolded (U)-like and native (N)-like, representing different extents of chain compaction (34, 35). The dependence of spatial separation on sequence separation indicated that the U-like sub-ensemble at 100 ms was random-coil-like with non-specific intra-chain interactions, while the N-like sub-ensemble had at least some specific native-like intra-chain contacts (34). Analysis of the kinetic evolution of the U-like to the N-like sub-ensemble for the B, C and E segments (35) led to a mechanism for chain contraction (Figure 1c), which closely resembled the previously established mechanism for folding (41). This suggested that, following the initial chain collapse at 100 ms, chain contraction and structure formation proceed concurrently during the folding of MNEI.

**Figure 1.**
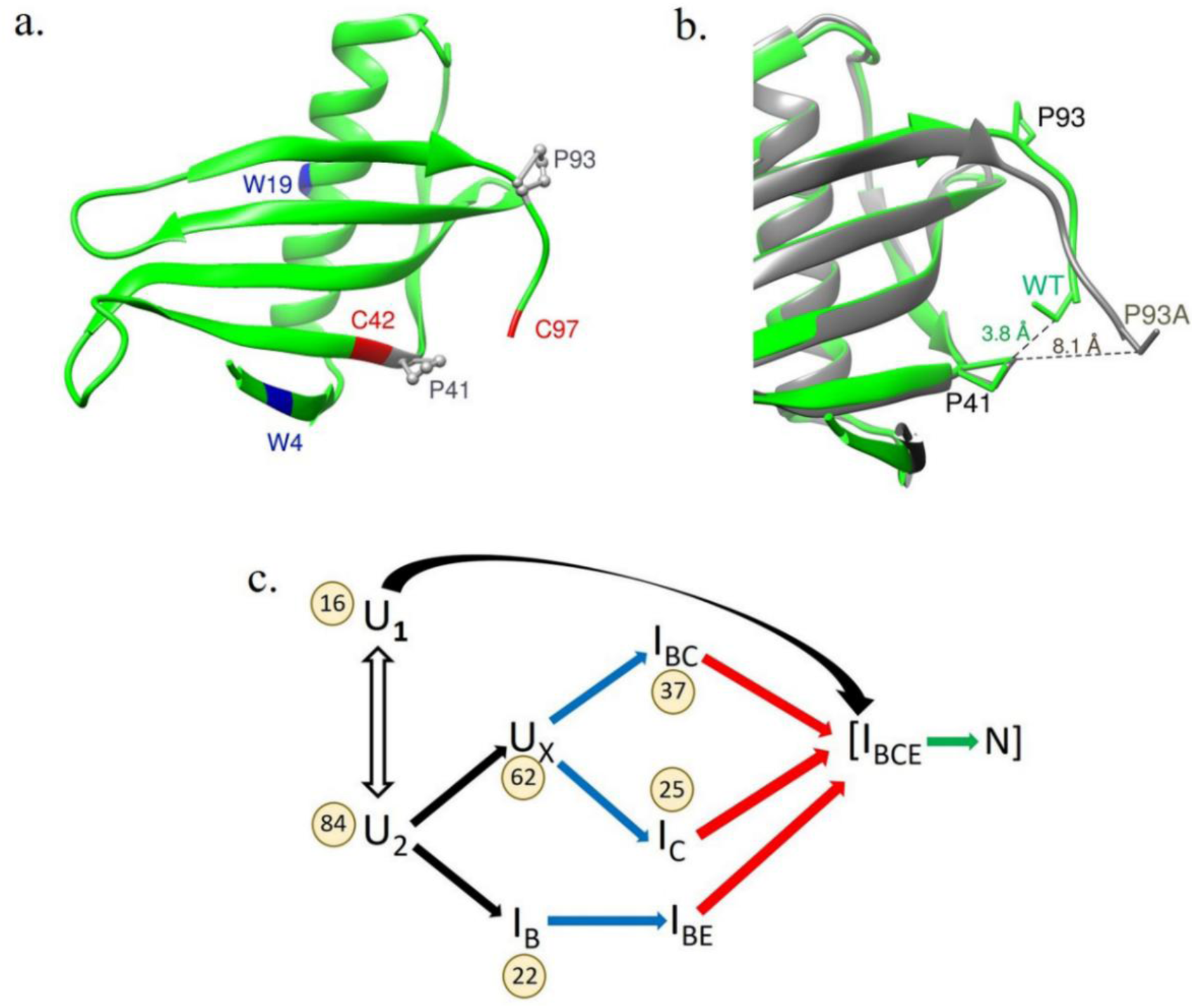
Structure and mechanism of folding/contraction of single-chain monellin (MNEI). a) The residues used for monitoring FRET, and the locations of the two native *cis*-Pro residues (Pro41 and Pro93, in gray) that were mutated to Ala, are shown. Trp4 and Trp19, which served as FRET donors, and Cys42 and Cys97, which served as FRET acceptors, are marked in blue and red, respectively. FRET was measured for the Trp19-Cys42TNB pair, for mapping the central core (C) segment, and for the Trp4-Cys97TNB pair, to map the end-to-end distance (E) segment. b) Structural superposition of WT MNEI (PDB ID: 1IV7) and its P93A mutant variant (PDB ID: 7EUA) highlighting differences in the C-terminus region, particularly its proximity to Pro41 in the β2 strand. The protein structures shown in panels a and b were drawn using Chimera. The distances between the side-chains are shown in Å. c) Scheme showing chain contraction during folding (35). The black, blue, red and green arrows represent the unobservable (complete within 100 ms), fast, slow and very slow kinetic phases of folding/contraction. The subscripts on the intermediate (I) denote the segments (B, C and/or E) which have contracted to N-like dimensions in that intermediate. The numbers in the circles indicate the percentage of molecules following a given pathway.

A key question is whether chain compaction can be separated temporally from structure formation steps at later stages of folding, just as initial chain compaction is decoupled from structure formation in the case of MNEI (28, 29) as well as other proteins (23–26). It is likely that sequence-encoded structural constraints such as those imposed by proline residues, modulate the coupling of chain contraction and structure formation not only during initial collapse but also at later stages of the folding reaction. In the case of MNEI, an earlier study had shown that mutation of the two proline residues (Pro41 and Pro93), which are *cis* in the N state, to alanine, simplified the folding landscape, and had indicated that chain contraction might also be modulated (38).

In the current study, site-specific Pro to Ala mutations were introduced at the two native-*cis* prolines (Pro41 and Pro93) of MNEI, and FRET pairs were designed to monitor compaction of distinct structural segments (Figure 1). The effect of the P41A mutation on compaction of the C segment was monitored using measurements of FRET between Trp19 and Cys42-TNB in both the WT protein (WT_C) and the P41A mutant variant (P41A_C) (Figure 1a). Similarly, the effect of the P93A mutation on compaction of the E segment was monitored using measurements of FRET between Trp4 and Cys97-TNB in both the WT protein (WT_E) and the P93A mutant variant (P93A_E). It should be noted that the FRET acceptor TNB is a small and charged moiety compared to the bulky and highly hydrophobic moieties (42, 43) that have been used as probes in other FRET studies. Because Pro41 and Pro93 are located in the proximity of the core (C segment) and the end-to-end (E segment) regions, respectively, their mutation has enabled site-specific probing of how proline residues influence local chain contraction at the C and E segments during folding. Distributions of intramolecular distances spanning the C and E segments and their contraction at different times of folding were characterized using MEM-coupled tr-FRET measurements. Folding itself was monitored using measurements of the changes in the fluorescence of Trp19 and Trp4 during the folding of the Trp19-containing and Trp4-containing protein variants, respectively. The measurements have provided mechanistic insight into the role of the proline residues in the contraction of protein regions in their proximity. Importantly, they have revealed that backbone rigidity can tune the coupling between chain compaction and structure formation during all stages of folding.

## Results and Discussion

### The structure and stability of MNEI are not affected significantly by Pro to Ala mutation and TNB-labeling

Far-UV circular dichroism spectra showed that both single Pro to Ala mutations, as well as TNB-labeling, had only small effects on the secondary structure of the protein (Figure S1). Fluorescence spectra showed that the environments of Trp19 and Trp4 remain largely unchanged upon Pro to Ala mutation (Figure S2). The fluorescence emission maxima of Trp19 in the N states of both WT_C and P41A_C were red-shifted compared to that of Trp4 in WT_E and P93A_E, indicating that the environment of Trp19 is more polar than that of Trp4 (44).

Upon TNB-labeling, the quenching of Trp19 fluorescence because of FRET across the C segment to the TNB adduct on Cys42, as well as the quenching of Trp4 fluorescence because of FRET across the E segment to the TNB adduct on Cys97, were both reduced upon Pro to Ala mutation. The fluorescence intensities of Trp19 and Trp4 in the N states of TNB-labeled P41A_C and P93A_E were 2.5-fold and 2-fold higher than those for TNB-labeled WT_C and WT_E, respectively (Figure S2). The FRET efficiencies calculated for the N states of WT_C and WT_E were 0.9 and 0.8, respectively, and decreased to 0.8 and 0.6 for the N states of P41A_C and P93A_E, respectively (Figure 2, insets). These observations suggested that the N states of P41A_C and P93A_E have slightly expanded C and E segments, respectively. In the case of P93A_E, this was consistent with the fact that the Pro93 to Ala mutation causes the C-terminal sequence segment 93-97 to move away from the protein core, because of the Gly92-Ala93 peptide bond adopting the *trans* conformation (Figure 1b).

**Figure 2.**
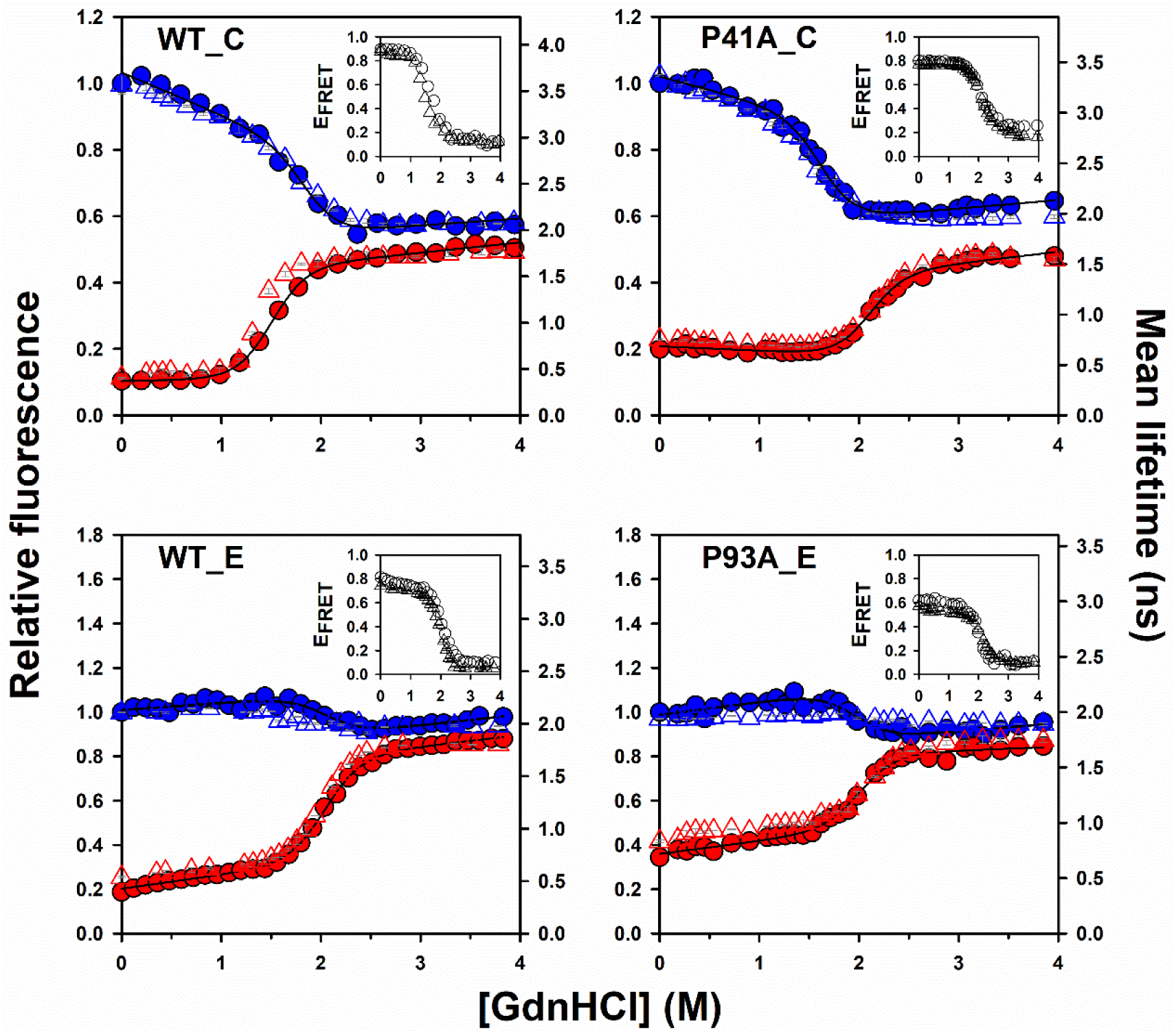
Equilibrium unfolding transitions of MNEI monitored by steady-state FRET (circles) and time-resolved FRET (triangles). Blue and red colors represent the fluorescence intensities and mean lifetimes of the unlabeled and TNB-labeled proteins, respectively. The solid lines through each dataset are non-linear least-squares fits to a two-state unfolding model (70). The thermodynamic parameters obtained are listed in Table S1. The inset in each panel shows the correspondence between FRET efficiency calculated from fluorescence intensity (circles) and fluorescence mean lifetime (triangles) measurements. The error bars represent the spread in the data obtained from two independent experiments. The data for WT_C and WT_E are from reference 44.

Neither the single Pro to Ala mutations nor TNB-labeling significantly affected the stability of the protein (Figure 2 and Table S1). P41A_C was marginally less stable than WT_C, and the stabilities of both proteins were not altered to significant extents upon TNB labelling (Table S1). TNB-labeling of Cys97 had no significant effect on the stabilities of WT_E and P93A_E. Overall, the effects of the mutations and TNB-labeling on the structure and dynamics were minor (Figures 2, S1 and S2). It should be noted that protein stabilities were determined by carrying out two-state analyses of the equilibrium unfolding transitions (44). It had been shown previously that fluorescence and CD-monitored equilibrium unfolding transitions yield the same values for the protein stability, for these four MNEI variants (38, 44), as well as for other MNEI variants (28, 29).

### Validation of the time-resolved fluorescence measurements

The equilibrium unfolding transitions monitored by measurement of the Trp fluorescence intensity at each GdnHCl concentration closely matched those monitored by measurement of the mean fluorescence lifetime determined from discrete analysis (see Materials and Methods, SI) of the Trp fluorescence decay measured at each GdnHCl concentration (Figure 2). This was an important result because it validated the accuracy of the fluorescence lifetime decay measurements and set the stage for analysis of the fluorescence decays by the maximum entropy method (MEM).

### MEM-analyzed fluorescence decays distinguish quantitatively between U-like and N-like sub-populations present together

MEM analysis (see Materials and Methods, SI) was used to convert fluorescence decays collected at different GdnHCl concentrations to fluorescence lifetime distributions for both the unlabeled (Figure S3) and labeled (Figure S4) protein variants. The MEM-derived distributions for all the unlabeled variants were largely unimodal (Figure S3). In contrast, those for the labeled variants were typically bimodal (Figure S4), and displayed a peak for lifetimes shorter than 0.6 ns, originating from a N-like sub-population (more FRET, hence shorter lifetimes), as well as a peak for lifetimes longer than 0.6 ns, originating from an U-like sub-population (less FRET, hence longer lifetimes) (3, 34). This assignment was based on the observation that the U state predominantly populates lifetimes > 0.6 ns (U-like) and the N state predominantly populates lifetimes < 0.6 ns (N-like) (Figure S4) (33, 35).

Such groupings of lifetimes allowed the fractions of molecules in the U-like and N-like sub-populations to be determined at each GdnHCl concentration. To extract these fractions from the MEM distributions, it was first necessary to account for the observation that for any segment, the equilibrium U state contained a fraction of molecules with lifetimes corresponding to the N-like distribution (< 0.6 ns), while the equilibrium N state contained a fraction of molecules with lifetimes corresponding to a U-like distribution (> 0.6 ns) (Figures S3 and S4). These fractions were comparable for each pair of unlabeled (Figure S3) and corresponding labeled (Figure S4) unfolded proteins, indicating that they originated from differences in the electronic structure of the fluorophore or from distinct Trp rotamers (45, 46), and not from the presence of the quenching TNB moiety. The fraction of molecules expanded (U-like) at a given segment was determined using the equation: f_U_=Y_i_−Y_N_/Y_U_−Y_N_, where Y_U_ represents the relative sum of amplitudes for the U-like distribution in the equilibrium U state, Y_N_ corresponds to the relative sum of amplitudes for the U-like distribution in the equilibrium N state, and Y_i_ denotes the relative sum of amplitudes for the U-like distribution at a given GdnHCl concentration. The relative sum of amplitudes was calculated as the sum of amplitudes for the U-like or N-like distributions divided by the total sum of amplitudes for both distributions. f_U_ derived from MEM-derived fluorescence lifetime distributions (Figure S4), had been shown previously to accurately estimate the relative fractions of N-like and U-like molecules present together (33, 35).

It should be noted that while the N states of WT_C, P41A_C, and WT_E, like those of several other proteins (44, 47, 48), displayed bimodal MEM-derived fluorescence lifetime distributions with a very minor (~10–20%) fraction of molecules appearing to be U-like (Figure S4), the N state of P93A_E exhibited a broad unimodal distribution with no clear distinction between the N-like and U-like sub-ensembles. Clearly, the Pro93 to Ala mutation made the N state more heterogeneous, with molecules exhibiting a broad range of end-to-end distances. It is possible that the *cis* Pro93 introduces a kink or bend in the protein backbone in the N state (Figure 1b), which may contribute to the stability of a specific conformation or sub-ensemble. Thus, in WT_E-TNB, two distinct conformations (N-like and U-like) might be stabilized, leading to a bimodal distribution. When Pro93 was mutated to Ala, a more flexible amino acid, the rigidity that Pro93 provided was lost. This increased flexibility likely allows the protein to sample a broader range of conformations, thereby blurring the separation between the two sub-ensembles, and resulting in the broad unimodal distribution seen for P93A_E (Figure S4). Importantly, although the distribution was unimodal, the relative sum of amplitudes for lifetimes >0.6 ns (U-like) was still ~25–30%, comparable to WT_E. Normalization of the amplitudes (see above) ensured that f_U_ could still be accurately determined.

For each protein variant, the dependence of the MEM-derived fraction of molecules that were U-like calculated as described above (Figure S5) was found to match the unfolding transition monitored by the measurement of fluorescence intensity or mean fluorescence lifetime of Trp4 or Trp19, and to yield similar values for the stability, ΔG_U_ and the midpoint of the transition, C_m_ (Table S1). This correspondence validates the use of the MEM-derived fluorescence lifetime distributions for determining accurately the relative fractions of N-like and U-like molecules present together, using the cut-off value of 0.6 ns for the fluorescence lifetime (see above).

### The Pro41 to Ala mutation stabilizes N-like structure at the C segment in the product of the initial collapse reaction

Transformation of the kinetic data in the insets of Figures S6 and S7 into FRET efficiency (Figure 3; see Materials and Methods, SI) revealed a distinct burst phase for both WT_C and P41A_C, which was more pronounced in the latter. One explanation for the observed change in FRET efficiency measured in this way could be that all molecules became compact initially, but that the extent of compaction was more in the case of P41A_C than for WT_C. In other words, it could be that the Pro41 to Ala mutation led to a greater reduction in the mean intramolecular distance at the C segment (insets, Figure 3). The alternative explanation could be that all molecules did not become compact, but only a sub-population of them underwent significant compaction, while another sub-population underwent little if any compaction. To distinguish between these possibilities and obtain quantitative insight into the fraction of molecules undergoing compaction, the fluorescence decays were analyzed using MEM (Figures S6, S7 and 4; see Materials and Methods, SI). As in the case of the equilibrium unfolding data (Figures S3 and S4), N-like and U-like sub-populations were observed at different times of folding. The kinetics of the decrease in the fraction of U-like molecules, f_U_ (described above), as they contracted to become N-like during folding, was determined (Figure 5).

**Figure 3.**
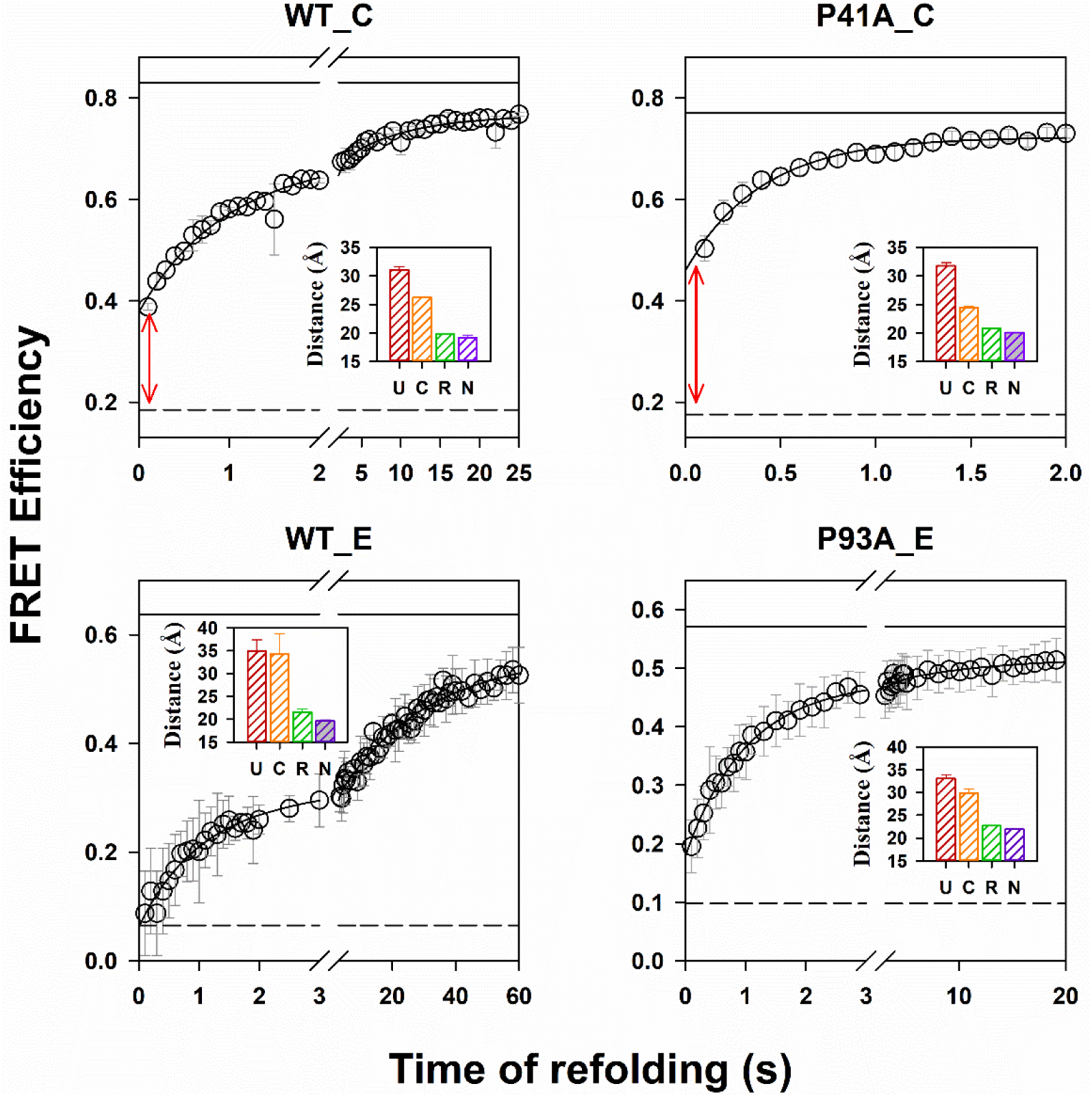
Kinetics of folding in 0.4 M GdnHCl monitored by trFRET for the different mutant variants of MNEI. FRET efficiency was calculated using the values of the mean lifetime of the unlabeled and TNB-labeled variants at each folding time point (equation 5, see Materials and Methods, SI). In each panel, the solid and dashed black horizontal lines represent the FRET efficiencies in the N and U states, respectively. The red arrow in each panel indicates the burst-phase change in FRET efficiency from the U state to the first observable folding time point (100 ms). The data for WT_C, WT_E and P93A_E were fit to a two-exponential equation, while that for P41A_C was fit to a single exponential equation. The rate constants and relative amplitudes obtained from the fitting are reported in Table 1. The inset in each panel shows the average donor−acceptor pair distance (<R_DA_>) calculated at different stages of folding (U, unfolded; C, collapsed intermediate at 100 ms; R, refolded; and N, native state) using FRET efficiency values and the Förster equation (equation 6, see Materials and Methods, SI). Error bars indicate standard errors from two independent double-kinetics experiments. Panels WT_C and WT_E have been adapted and modified from reference 35.

**Table 1.** Rate constants and their corresponding relative amplitudes obtained by fitting the kinetic phases of folding/contraction.

| Protein | Phase | Rate constants (amplitudes) of folding<sup>a</sup> (s<sup>-1</sup>) | Rate constants (amplitudes) of the ss-FRET efficiency monitored change<sup>b</sup> (s<sup>-1</sup>) | Rate constants (amplitudes) of the tr-FRET efficiency monitored change<sup>c</sup> (s<sup>-1</sup>) | Rate constants (amplitudes) of the decrease in U-like population<sup>d</sup> (s<sup>-1</sup>) |
| --- | --- | --- | --- | --- | --- |
| WT_C | Burst | — | — (27%) | — (39%) | — (16%) |
|  | Very fast | 4.7 (14%) | 7.5 (19%) | — | — |
|  | Fast | 0.6 (60%) | 1 (40%) | 1 (43%) | 2 (61%) |
|  | Slow | 0.06 (26%) | 0.07 (14%) | 0.1 (18%) | 0.17 (23%) |
| P41A_C | Burst | — | — (44%) | — (58%) | — (63%) |
|  | Very fast | 10.5 (12%) | 34 (14%) | — | — |
|  | Fast | 1.5 (64%) | 2.8 (38%) | 2 (42%) | 2 (37%) |
|  | Slow | 0.14 (24%) | 0.16 (4%) | — | — |
| WT_E | Burst | — | — | — | — (11%) |
|  | Very fast | 25 (25%) | 26 (21%) | — | — |
|  | Fast | 1 (48%) | 1 (26%) | 1 (42%) | 2 (19%) |
|  | Slow | 0.1 (27%) | 0.03 (53%) | 0.03 (58%) | 0.04 (70%) |
| P93A_E | Burst | — | — (14%) | — (17%) | — (15%) |
|  | Very fast | — | — | — | — |
|  | Fast | 1 (78%) | 1 (70%) | 1 (69%) | 1 (75%) |
|  | Slow | 0.1 (22%) | 0.1 (16%) | 0.1 (14%) | 0.1 (10%) |

<sup>a</sup> Measured from the kinetics of change in the fluorescence of Trp19 or Trp4 in the unlabeled proteins.
<sup>b</sup> Measured from the kinetics of change in fluorescence intensity of Trp19 or Trp4 in the TNB-labeled and unlabeled proteins. The burst phase was of 10 ms duration.
<sup>c</sup> Measured from the kinetics of change in the mean fluorescence lifetime of Trp19 or Trp4 in the TNB-labeled and unlabeled proteins, shown in Figure 3. The burst phase was of 100 ms duration.
<sup>d</sup> Measured from the kinetics of change in the U-like population seen in the MEM distributions of TNB-labeled proteins, shown in Figure 5. The burst phase was of 100 ms duration.
The data for WT\_C and WT\_E have been taken from reference 35. The errors in the rate constants and their corresponding relative amplitudes are within $\pm 10\%$ .

The major effect of the P41A mutation was that the fraction of molecules that become N-like at the C segment at 100 ms of folding, increased from 16 % in the case of WT_C to 63 % for P41A_C (Figure 5, Table 1). It is important to explain this observation in the context of the mechanism of chain contraction and folding proposed for the wild-type protein (Figure 1c), for which the collapsed intermediate ensemble populated at 100 ms consists of three sub-ensembles (Figure 1c). 16% are present as I_BCE_ in which segments B, C, and E have all collapsed to become N-like; 22% are present as I_B_ in which only segment B has collapsed to become N-like; and 62% remain U-like, as U_X_, at all three segments. It appears that the P41A mutation could have resulted in two different types of effects on the heterogeneity of this collapsed ensemble formed at 100 milliseconds of folding. (1) It could have led to the induction of contraction of segment C in some of the U_X_ and I_B_ molecules. (2) It could have led to the conformational selection of very minor pre-existing sub-ensembles present in equilibrium with U_X_ and I_B_, in which the C segment is already N-like in its dimensions.

The P41A mutation does not affect the heterogeneity in the unfolded state (38); hence, the 63% molecules that have collapsed at 100 ms in the case of P41A_C would include the 16% which folded during the very fast phase (originating from U_1_ sub-ensemble) leading to the formation of I_BCE_ (Figure 1c). Since the slow phase of contraction at the C segment was absent for P41A_C (Figure 5, Table 1), it seems likely that the 22% molecules that contract slowly in the case of WT_C (Figure 1c), instead collapse during the burst phase on the same pathway. To account for the remaining 25% (63-(16+22)) molecules that collapsed at the C segment during the burst phase in the case of P41A_C, it is necessary to posit that the mechanism of folding/contraction for WT_C (Figure 1c) has been altered by the P41A mutation to that shown below in Scheme 1.

**Scheme 1.**
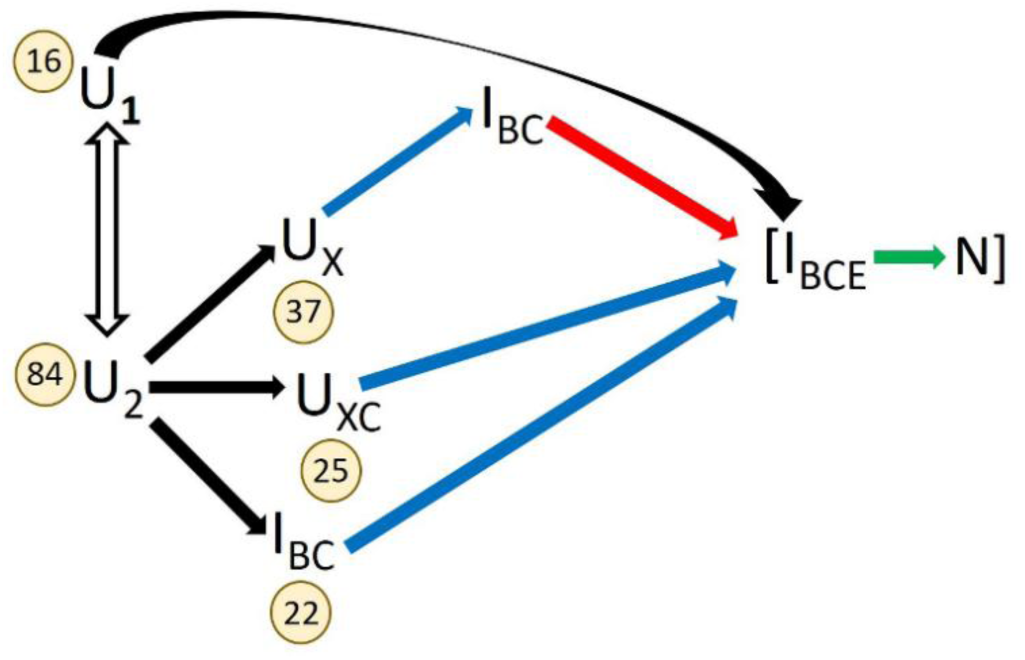

Several salient features of Scheme 1 need to be noted. (1) In the case of WT MNEI (WT_C), the kinetic data had indicated that the 100 ms burst phase led to the formation of two burst phase intermediates, U_X_ and I_B_, from U_2_, and that intermediates I_C_ and I_BC_ formed by kinetic partitioning from U_X_ (Figure 1c) (35). It seems now, that U_X_ is not homogeneous, but consists of two sub-populations U_X_ and U_XC_. In the case of WT_C, U_XC_ is populated to a negligible, undetectable extent, but in the case of P41A_C, it becomes populated to 25%. It appears that the P41A mutation significantly stabilizes U_XC_ by relieving the conformational strain imposed by Pro41. Stabilization may also occur *via* an intramolecular hydrogen bond, in which the introduced Ala, unlike Pro, can participate; such bonds have been suggested to facilitate initial polypeptide chain collapse (49–52). (2) The stabilization afforded by the mutation is not large, and 37% molecules remain as U_X_ at the end of the burst phase. In these molecules, the C segment becomes N-like only at the end of the fast phase of folding, along with the B segment. Hence, the fast phase of contraction at the C segment seen for WT_C is also seen for P41A_C. (3) The C segment, which was U-like in I_B_ in the case of WT_C, is stabilized by the P41A mutation so that it becomes N-like in the case of P41A_C; hence, I_B_ is shown as I_BC_. Consequently, the slow phase of contraction at the C segment seen for WT_C (Figure 1c) was not seen for P41A_C. The effects of the P41A mutation described above are consistent with known effects of mutations in modulating the kinetic partitioning between competing folding pathways by stabilizing or destabilizing specific intermediates (53–56). (4) It is not known whether the P41A mutation has a similar effect on the B segment (β-sheet). It will be important to investigate this in the future, especially since residues in the β-sheet contribute to the protein core, and since studies with other proteins have reported coupling o β-sheet formation with hydrophobic core packing (57, 58). It is therefore not surprising that the β-sheet and core both become N-like together on the U_2_ → I_BC_ → I_BCE_ and U_2_ → U_X_ → I_BC_ → I_BCE_ pathways.

### The Pro93 to Ala mutation stabilizes N-like structure at the E segment in the products of the fast phase of collapse

In the case of WT_E, only 22% of the molecules were found to contract their end-to-end distance (the E segment) during the fast phase of folding when they form I_BE_ (Figure 1c). The contraction of the E segment in the remaining molecules was found to occur primarily during the slow phase (Figures 1c and 5; Table 1). However, in the case of P93A_E, 75 % molecules were found to contract during the fast phase, and only about 10% did so in the slow phase. Hence, the P93A mutation leads to the contraction of an additional 53% (75-22) of molecules at the E segment to become N-like during the fast phase of contraction. To account for these additional 53% molecules, it becomes necessary to propose that I_BC_ and I_C_ (Figure 1c) also exist in equilibrium with a so far hidden I_BCE_ and a hidden I_CE_, respectively. Because of the strain introduced in the polypeptide backbone by Pro93, I_BCE_ and I_CE_ are unstable and hence populated to an insignificant extent in the case of WT_E. In the case of P93A_E, they became stabilized and populated to a larger extent because of the replacement of Pro93 with Ala. The P93A mutation therefore leads to conformational selection of I_BCE_ and I_CE_ such that they are together populated to about 53%, and I_BC_ and I_C_ are consequently together populated to only about 10 %. Thus, the 75 % molecules contracting their E segment in the fast phase in the case of P93A_E, are those which form I_BE_, I_CE_ and I_BCE_.

It should be noted that *trans* to *cis* isomerization of Pro93 occurs during the slow phase of folding of WT_E (38). In this context, an alternative explanation could have been that the contraction that occurs during the slow phase of folding of WT_E is only that which accompanies the *trans* to *cis* isomerization of the Gly92-Pro93 bond, which brings the FRET acceptor, Cys97TNB as well as the residues 93-97 closer to the FRET donor, Trp4 and the core of the protein (Figure 1b). In the case of P93A_E, the Gly92-Ala93 bond remains *trans*, and does not undergo *trans* to *cis* isomerization; consequently, the FRET acceptor does not move towards the donor (Figure 1b), and no slow phase contraction would be observed. While this explanation would have been difficult to rule out if only FRET measurements had been carried out, it can be ruled out now based on the MEM analysis which showed that about 10% molecules still contract during the slow phase in the case of P93A_E (Figure 5, Table 1).

Thus, the result that the P93A mutation alters the population of intermediates by stabilizing the E segment in I_CE_ and I_BCE_, which were populated only insignificantly in the case of WT_E, mirrors the effects observed for the P41A mutation (see above). This suggests that Pro41 and Pro93 dictate the partitioning of pathways, and play a crucial role in determining the stability of specific intermediates and hence, the heterogeneity of the intermediate ensemble. Similar proline-mediated modulation of pathway partitioning has also been observed for other proteins (59–61), but has not been directly quantified. This is perhaps the first experimental study to directly quantify the redistribution of folding intermediates arising from the gain in backbone flexibility consequent to replacing rigid Pro with flexible Ala. By quantitatively tracking shifts in the distribution of coexisting sub-populations within intermediate ensembles, this study provides direct evidence that the stability and partitioning of sub-populations are tunable features encoded in the primary sequence.

In summary, the Pro41 to Ala mutation has revealed hidden heterogeneity in the size of the C segment in the product of initial collapse. Similarly, the Pro93 to Ala mutation has revealed hidden heterogeneity in the size of the E segment in a late intermediate. In previous studies of the folding of barstar, it had been shown that the product of initial collapse was also heterogenous, with different structural segments collapsed to different extents (11, 27). It was shown that this collapsed state at a few milliseconds of folding also contained sub-populations of differently structured molecules, and that different structured sub-populations could become stabilized and hence be populated to detectable extents in different folding conditions (62, 63). Previous trFRET studies of the slow folding of barstar had shown that the structural heterogeneity of a late folding intermediate could also be modulated by a change in folding conditions (64). The current study demonstrates that structural heterogeneity in intermediate ensembles can be modulated by site-specific mutations that relieve local backbone rigidity, thereby redistributing molecules among distinct sub-populations (see above). Consequently, backbone rigidity emerges as a key determinant of how structural heterogeneity is modulated during protein folding.

### Decoupling of structure formation and chain compaction during folding

Different segments of the polypeptide chain of WT MNEI were shown to collapse to different extents during the burst phase of folding, suggesting that the product of the burst phase is structurally heterogeneous (28). It was subsequently determined that the product of folding at 100 ms is also heterogeneous, comprising sub-populations of molecules that differed in which of the B, C and E segments had become N-like in dimensions (35), even though insignificant secondary structure has formed at this time (28, 29, 33). These results indicated that initial collapse at 100 ms of folding is decoupled from structure formation. In this study, it is seen that the Pro41 to Ala mutation further decouples initial collapse at the C segment from structure formation, by increasing the number of molecules becoming N-like in their dimensions during initial collapse (Figure 5). This increase in decoupling because of the P41A mutation suggests that rigid proline residues may serve as local gatekeepers of the initial collapse process. While initial collapse may be hydrophobic in nature, it also appears to be under precise, residue-level control.

Importantly, the increase in the number of molecules undergoing initial collapse occurred at the expense of fewer molecules contracting during the subsequent phases, even though the number of molecules undergoing a change in structure in each phase, measured by the change in fluorescence of Trp19, was affected only marginally (Table 1). In this context, it should be noted that the rate constants of the change in fluorescence were found to match those of the change in far-UV circular dichroism, for WT_C and WT_E, as well as for other MNEI variants (28). Similarly, it was also seen that the Pro93 to Ala mutation increases the fraction of molecules becoming N-like in dimensions during the fast phase of contraction, at the expense of the slow phase (Figure 5) even though the number of molecules undergoing a change in structure in each phase, was affected only marginally (Table 1). The very fast phase of folding observed for WT_E (~25% of molecules) is abolished in the case of P93A_E (38), with these molecules now folding during the fast phase. Thus, while the overall number of molecules folding remains unchanged, the extent of contraction during the fast phase is clearly increased. Thus, it appears that while an intermediate ensemble may consist of sub-populations that have compacted differently in different segments, the sub-populations nevertheless do not differ in the extent of structure that has formed (schematically illustrated in Figure S8). Together, these results show that the Pro to Ala mutations lead to more contracted sub-populations being favored, without significantly altering the extent of structure formed. This decoupling of chain contraction from structure formation demonstrates that the two processes, though often concurrent, are separable events during folding.

Local backbone rigidity therefore appears to modulate the extent of compaction occurring at different stages of folding, without affecting the extent of structure formed. These effects, although mechanistically different for the two Pro to Ala mutations, demonstrate collectively that structure formation can be decoupled from chain contraction during all kinetic phases of folding, just as it is known to be decoupled during the initial collapse reaction. This appears to be first study that directly demonstrates the decoupling of chain contraction and structure formation during the later stages of folding. The results reveal that folding occurs through separable physical events, chain compaction and structure formation, which are not obligatorily coupled to each other, even at later stages of folding.

### Contraction occurs gradually during folding and is accelerated because of Pro to Ala mutations

The trFRET measurements showed that the U-like sub-population contracts to an N-like sub-population during the fast and slow phases of folding, as well as during an unobservable 100 millisecond burst phase in the case of WT_C and WT_E (Figure 5, Table 1). These measurements also showed that each kinetic phase does not represent a two-state transition but is characterized by molecules contracting in a gradual manner (Figure 6). In the case of WT_C and WT_E, both the N-like and U-like sub-populations underwent gradual contraction during folding, as evident in the gradual shift in their MEM-derived distance distributions, and their peaks, toward shorter distances (Figures 4 and 6). An earlier report (34) attributed this contraction to continuous breakage of non-native interactions. Surprisingly, however, in the case of P41A_C, the N-like sub-population did not undergo any contraction during folding (Figure 6). It seems that the release of strain in the polypeptide backbone, and/or introduction of additional hydrogen bonding, consequent to replacing Pro41 with Ala, allowed the N-like sub-population to relax to the dimensions of the final N state as soon as it was formed. Moreover, the U-like sub-population for P41A_C undergoes contraction only during the fast phase (the slow phase was absent, see above and Figure 5). This indicates that compaction at the C segment is accelerated as a result of the Pro41 to Ala mutation. In the case of WT_E, the gradual nature of the slow compaction phase may be modulated by the *trans*-to-*cis* isomerization of the Gly92–Pro93 peptide bond. In the case of P93A_E, for which contraction was seen to occur only during the fast phase of folding, this contraction was also observed to be gradual in nature, and not to be all-or-none (Figure S9).

**Figure 4.**
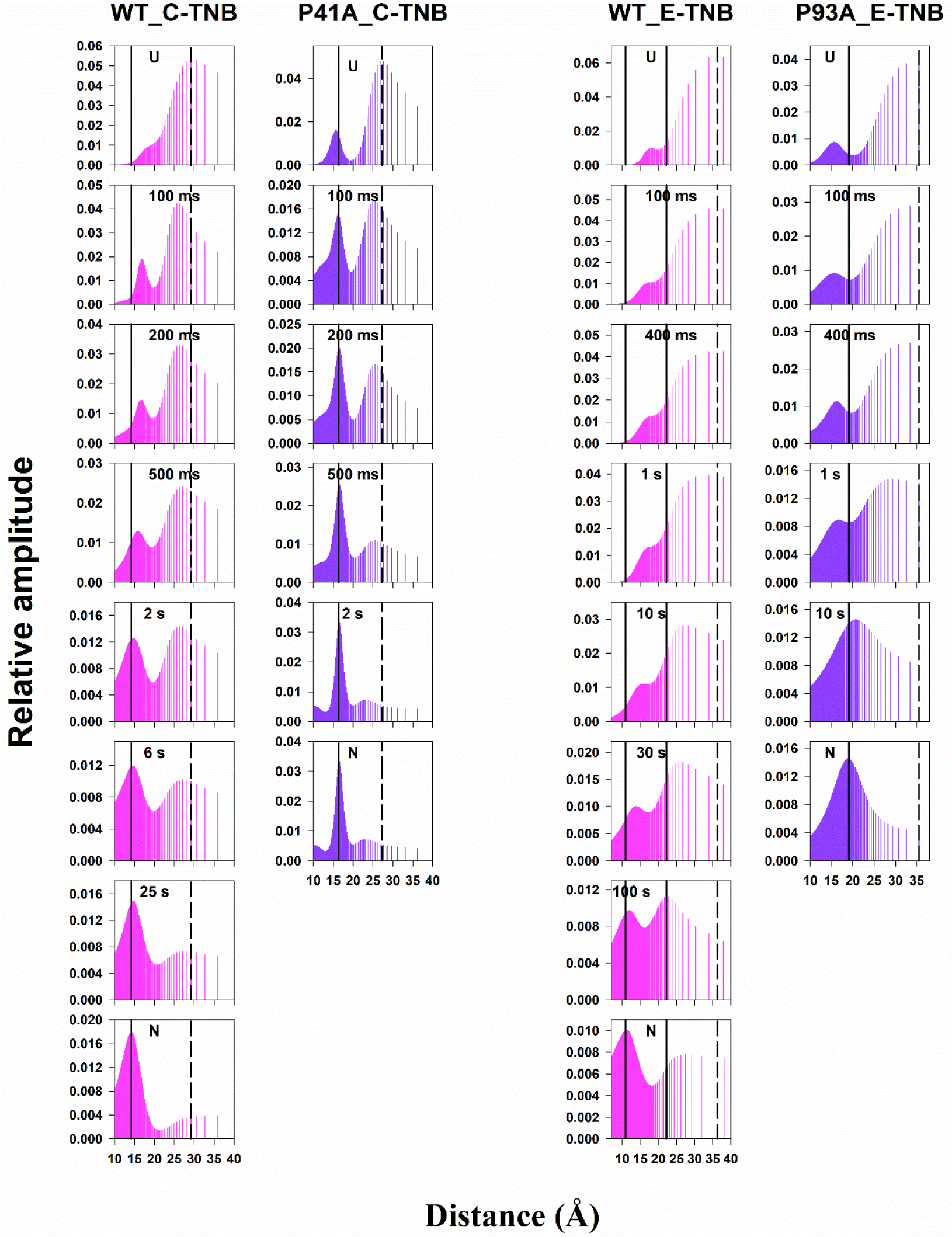
Evolution of distance distributions as a function of the time of folding following a 4 to 0.4 M GdnHCl jump. Experimentally derived distance distributions at representative time points of the folding reaction. The topmost panel is for the U state, the bottom-most panel corresponds to the distance distribution of the refolded N state, and the middle panels correspond to intermediate time points during folding as described in each panel. The vertical solid and dashed black lines indicate the peak positions of the distance distributions corresponding to the refolded N state and U state, respectively. Panels WT_C and WT_E have been adapted and modified from reference 34.

Nonetheless, for both Pro mutant variants, the U-like sub-population contracted gradually while at the same time undergoing first order transitions to become N-like, as also seen for the WT proteins (Figures 5 and 6). Similarly, gradual exposure of amide sites to solvent had been observed to occur in multiple kinetic phases in hydrogen exchange-mass spectrometry (HX-MS) studies of the unfolding of MNEI in native and native-like conditions (65, 66), in which folding would occur by the same mechanism in reverse. The HX-MS studies were complemented by SH-labeling studies which revealed that each kinetic pause during folding in native conditions occurs because a specific tight packing interaction has to form, and that interaction forms in an all-or none manner (67).

**Figure 5.**
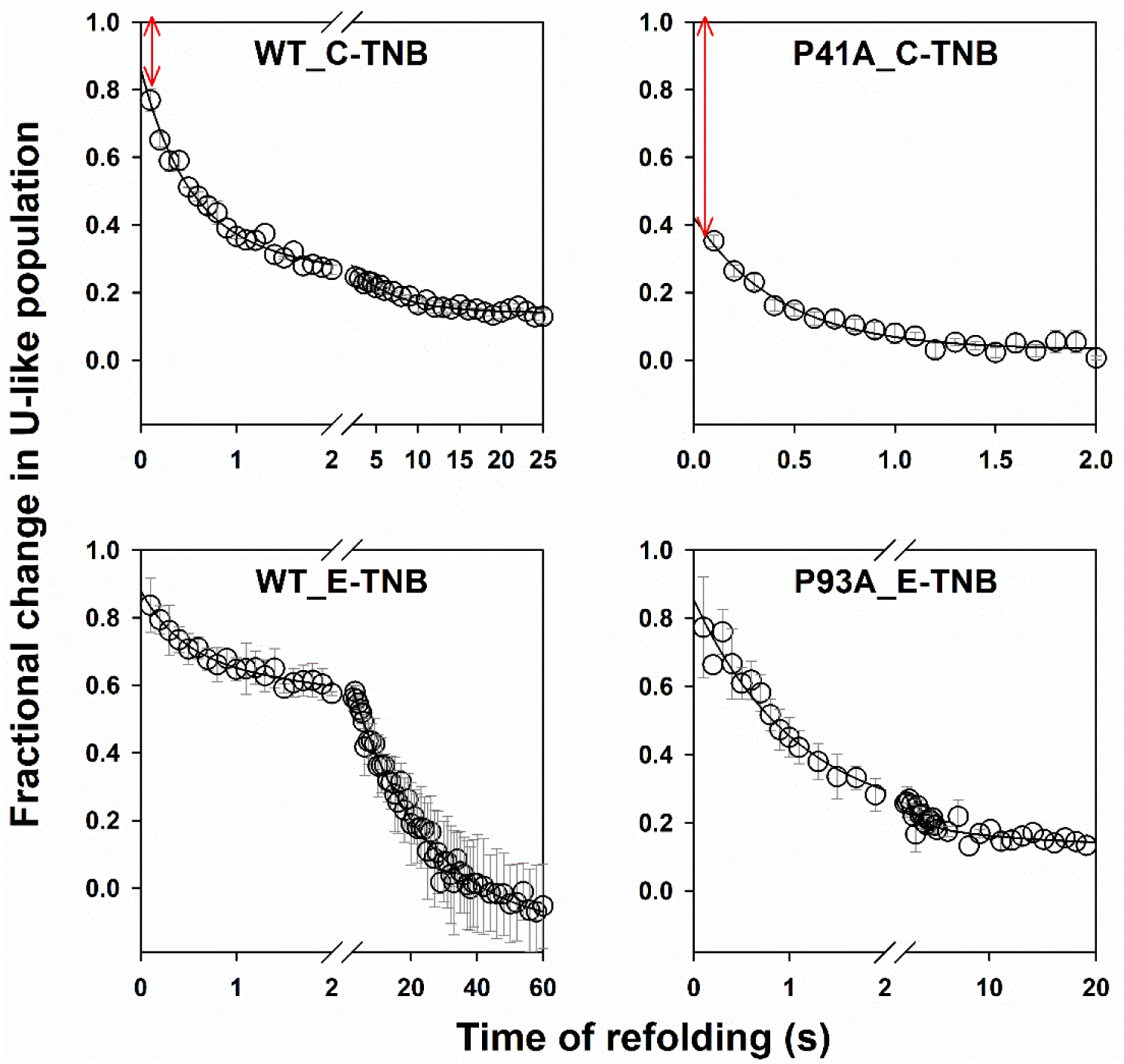
Kinetics of conversion from the expanded, U-like to the contracted, N-like population, measured at segments C and E for the different mutant variants of MNEI. The fractional population of molecules that are expanded (U-like) was determined as described in the text. The data for WT_C, WT_E and P93A_E were fit to a two-exponential equation, while that for P41A_C was fit to a single-exponential equation. The rate constants and relative amplitudes obtained from the fitting are reported in Table 1. The error bars represent the standard errors of measurements from at least two independent double kinetics experiments. Panels WT_C-TNB and WT_E-TNB have been adapted and modified from reference 35.

**Figure 6.**
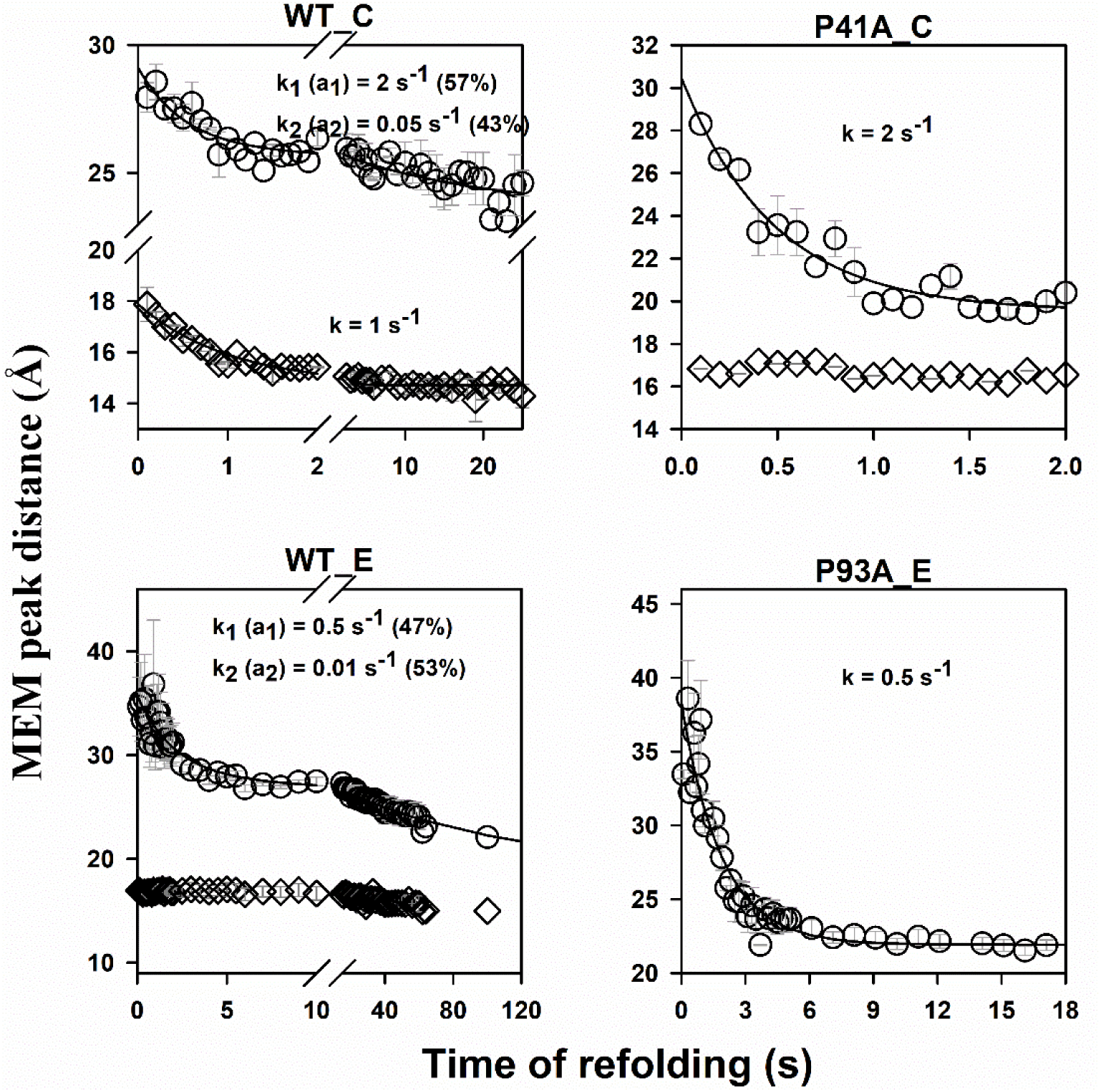
Kinetics of contraction of the U-like (circles) and N-like (diamonds) populations for different mutant variants of MNEI during folding in 0.4 M GdnHCl. For each mutant variant, the peak distance at each folding time was obtained using the MEM peak value of the U-like population and the N-like population in the MEM-derived fluorescence lifetime distribution obtained for the TNB-labeled variant (Figure S7) and the corresponding unlabeled counterpart (Figure S6) using equations 5 and 6 (see Materials and Methods, SI). The solid lines through the kinetic data represent fits to either a single or double exponential equation. The error bars represent the standard errors of measurements from at least two independent kinetic experiments. Panels WT_C and WT_E have been adapted and modified from reference 34.

The shift in the MEM peak position provides only qualitative insight into compaction (Figure 6). Acceleration of the kinetics of compaction is obvious in the case of the P41A_C, as a slow phase of compaction was not observed (Figure 5; Table 1). To quantify the mutation-induced acceleration of the compaction of the E segment during the fast and slow kinetic phases, the apparent time constants (determined as the amplitude-weighted average of the time constant, τ_av_ = (a_f_τ_f_ + a_s_τ_s_)/(a_f_ +a_s_); τ_i_ =1/k_i_) were calculated for WT_E and P93A_E, where τ_f_ and τ_s_ are the time constants of the fast and slow kinetic phases, and a_f_ and a_s_ are their respective relative amplitudes (Figure 5 and Table 1). The apparent time constants for WT_E and P93A_E were 20 s and 2 s, respectively, indicating a ten-fold acceleration in compaction due to the P93A mutation. The acceleration at both the C and E segments may arise, in part, from the replacement of the rigid Pro residue with a more flexible Ala, facilitating faster chain contraction, as previously observed (68, 69). Moreover, early stabilization of structure in both segments (see above) by Pro to Ala mutation, may pre-organize local and/or non-local conformations, leading to a reduction in the conformational degrees of freedom and enhancement in the efficiency of subsequent conformational search.

## Conclusion

Site-specific Pro to Ala mutations reveal how the heterogeneity within intermediate ensembles modulates the coupling between chain compaction and structure formation during the folding of MNEI. The P41A mutation stabilized a minor sub-population of molecules during the initial collapse process, while the P93A mutation stabilized late intermediates during the fast phase of compaction at the C and E segments, respectively. Stabilization of these sub-populations led to pathway partitioning and redistribution of intermediate sub-populations, resulting in a decoupling of structure formation from chain compaction during all kinetic phases of folding. These findings highlight a general principle: sub-population heterogeneity, encoded by local backbone rigidity, is a key determinant of whether and how compaction and structure formation are dynamically coupled during folding, challenging the traditional view that these processes are obligatorily coupled during the later stages of folding.

## ASSOCIATED CONTENT

### Supporting Information

SI contains Materials and Methods section, and Supplementary figures.

## Acknowledgements

We thank Drs S. Bhatia, M.K. Mathew and G. Krishnamoorthy, as well as members of our laboratory, for discussion, and Dr S. Bhatia for her comments on the manuscript. JBU is a recipient of a JC Bose National Fellowship from the Government of India. This work was funded by National Centre for Biological Sciences, Bengaluru, by the Indian Institute of Science Education and Research Pune, as well as by a grant (JBR/2021/000029) from the Science and Engineering Board, Government of India.

