## Supplementary Information for "Intermediate heterogeneity modulates coupling between chain compaction and structure formation during protein folding"

#### Materials and Methods

In this study, intramolecular FRET changes at two specific segments of MNEI were investigated. For FRET measurements, a Trp residue served as the donor fluorophore, while a Cys residue was covalently linked to the thionitrobenzoate (TNB) moiety, acting as the FRET acceptor. The two segments studied were the core (C) segment, spanned by Trp19-Cys42TNB FRET pair, and the end-to-end (E) segment, spanned by Trp4-Cys97TNB FRET pair. The effect of the Pro41 to Ala mutation on changes in compaction of segment C and the effect of Pro93 to Ala mutation on changes in the end-to-end distance (segment E) were studied. Of the six Pro residues in MNEI, only Pro41 and Pro93 adopt *cis* conformations in the native (N) state (1).

For each FRET pair, fluorescence measurements were carried out on both the unlabeled and the corresponding TNB-labeled variants. FRET was assessed using both steady-state and time-resolved measurements. In the case of steady-state FRET measurements, Trp fluorescence was measured in the absence (donor only) and presence of the acceptor moiety, TNB (donor–acceptor), with a dead time of approximately 10 ms. For time-resolved FRET measurements, fluorescence decay curves were collected either as a function of GdnHCl concentration or as a function of folding time using a double-kinetics setup, both in the absence and presence of the FRET acceptor. Because fluorescence intensity decays occur on a nanosecond timescale, much faster than the timescale of conformational fluctuations, time-resolved FRET enables monitoring of changes in the distribution of donor–acceptor distances within the population of molecules as a function of folding time, as demonstrated previously (2, 3). Information on FRET efficiency, average intramolecular distances, and fluorescence lifetime distributions was extracted from the decay curves using established methods (4, 5).

#### Proteins and Reagents

The expression, purification, and labeling of the mutant variants used in this study have been described previously (1, 3). All the experiments were carried out at pH 8.0 and 25 °C. The details of the reagents used in the experiments have been reported earlier (3).

**Site-directed Mutagenesis**

The Pro41 to Ala mutation was introduced in the mutant variant W19C42, as described in a previous study (1). The Pro93 to Ala mutation was introduced in the mutant variant W4C97, using the QuickChange site-directed mutagenesis method developed by Stratagene.

**Measurement of fluorescence and far UV CD spectra**

Fluorescence spectra were measured using a Fluoromax 4 (Horiba) spectrofluorometer. Protein samples were excited at 295 nm, and emission spectra were recorded from 305 to 450 nm with a response time of 1 s. The excitation and emission slit widths were set to 1 nm and 5 nm, respectively. The protein concentration was 10 μM. Each spectrum represents the average of three independent emission scans.

Far-UV circular dichroism (CD) spectra were obtained using a Jasco J815 spectropolarimeter and a cuvette with a 0.1 cm path length. Data were collected with a 1 nm bandwidth, a response time of 1 s, and a scan speed of 20 nm $min^{-1}$. The protein concentration was 10 μM. Each spectrum was generated by averaging twenty individual wavelength scans.

**Steady-state and time resolved FRET-monitored equilibrium unfolding experiments**

For GdnHCl-induced equilibrium unfolding experiments, 10 mM of protein was incubated for >6 h in different concentrations of GdnHCl (0–4 M). The equilibrated unlabeled and labeled protein samples were excited at 295 nm, and the fluorescence signals were monitored on an MOS-450 optical system (Biologic), using a 360 ± 10 nm band-pass filter (Asahi spectra) for steady-state FRET measurements. For time-resolved FRET measurements, fluorescence lifetime decay curves were acquired for both the unlabeled and labeled proteins using the time correlated single photon counting system set-up as described in a previous study (6). A two-state, N ↔ U model was used to fit the sigmoidal equilibrium unfolding transitions (7) to obtain the values for the free energy of unfolding in water, $\Delta G_U$, and the midpoint of transition (Cm).

**Trp-fluorescence monitored folding kinetics experiments**

The folding reaction was initiated by rapidly mixing unfolded protein (in 4 M GdnHCl) with folding buffer (containing no GdnHCl) in a 1:9 ratio using a stopped-flow cuvette (FC20) and a stopped-flow module from Biologic (SFM-400). The protein concentration was 10 μM. The dead time of stopped-flow mixing was ~10 ms.

**Double-kinetics experiment**

Simultaneous acquisition of fluorescence decay traces and fluorescence intensities was performed at different time points (every 100 ms) during the folding reaction, using a pulsed laser (Mai Tai HP, Spectra-Physics) as the excitation light source (2). Notably, the simultaneous acquisition of steady-state fluorescence intensity served to validate the accuracy of the double-kinetics data for each mixing experiment (Figures S6 and S7). Photon arrival data were collected using a single-photon counting card module (SPC-630) from Becker and Hickl. A detailed description of the double-kinetics setup and fluorescence data collection has been reported previously (2). The final protein concentration used was ~20 μM.

**Analysis of the fluorescence lifetime decay traces:** Details of the analysis are given in an earlier study (6). A brief description is given below:

**Discrete analysis:** The decay traces were fit to a sum of exponentials,

$$F(t) = \sum_{i=1}^{n} \alpha_i e^{-\left(\frac{t}{\tau_i}\right)} \quad \textbf{(1)}$$

Here, $\alpha_i$ is the relative amplitude of the $\tau_i$ lifetime component, t is time and n ranges from 2 to 3.

An amplitude-weighted mean lifetime, the mean lifetime, $\boldsymbol{\tau}_m$ at every time point of the refolding reaction was determined, was caculated as:

$$\boldsymbol{\tau}_m = \frac{\sum \alpha_i \tau_i}{\sum \alpha_i}; \sum a_i = 1 \qquad \textbf{(2)}$$

**MEM analysis:** The analysis begins with the assumption that the decay corresponds to a distribution of 100-150 lifetimes in the range 10 ps to 10 ns. The a priori distribution of lifetimes was assumed to be uniform in the logarithms of lifetimes being uniformly distributed in this range. Then, the best fit values of $\alpha_i$ and $\tau_i$ are determined using the Maximum Entropy Method (MEM) (3, 5).

The a posteriori distribution of these parameters was obtained by maximizing the Shannon Jaynes entropy S, defined as

$$S = -\Sigma\, P_i \log(P_i) \qquad \textbf{(3)}$$

$P_i$ is the normalized amplitude of the ith lifetime.

$$P_i = \frac{\alpha_i}{\sum \alpha_i} \qquad \textbf{(4)}$$

$\alpha_i$ is the amplitude of the ith lifetime.

Analysis parameters were optimized for obtaining precise and reproducible MEM distributions. The most probable (MEM peak) lifetime refers to the lifetime corresponding to the maximum amplitude in the lifetime distribution data.

**FRET efficiency and distance determination**

The mean FRET efficiency (<$E_{FRET}$>) was obtained from mean fluorescence lifetimes for the unlabeled (<$\tau_D$>) and the corresponding TNB-labeled (<$\tau_{DA}$>) variants using the following equation:

$$<E_{FRET}> = 1 - \frac{<\tau_{DA}>}{<\tau_D>} \quad \textbf{(5)}$$

The mean fluorescence lifetimes were determined by discrete analysis of the fluorescence decay traces as described earlier (6).

The average FRET efficiency values were converted to average intramolecular distance (<$R_{DA}$>) using the Förster equation:

$$<R_{DA}> = R_0 \left(\frac{1 - <E_{FRET}>}{<E_{FRET}>}\right)^{1/6} \quad \textbf{(6)}$$

The values for the Forster's distance, $R_0$ used for the WT proteins and their determination has been described in an earlier study (6). Upon Pro mutation to Ala, the $R_0$ values remained similar to those of their respective background constructs (24.5 Å for P41A and 22.7 Å for P93A).

**Converting fluorescence lifetime distributions into distance distributions**

Fluorescence lifetime distributions of both the unlabeled and the TNB labeled variants were determined for consecutive acquisition time windows of 100 ms by fitting the observed fluorescence decay profiles to the MEM algorithm (described above). The lifetime distributions of the TNB-labeled variant at different times of folding were then converted to FRET efficiency distributions by using the peak lifetime values obtained for the corresponding unlabeled variant (see Eq. 5 above). The FRET efficiency distributions were then transformed into distance distributions using the well-known Forster relation (see Eq. 6 above), as established in previous studies (2, 5). It should be noted that the cut-off of 0.6 ns used to distinguish between the Nlike and U-like subensembles in the fluorescence lifetime distributions corresponds to a distance cut-off of ~ 20 Å in the distance distributions, as calculated using equations 5 and 6.

### Supplementary Figures

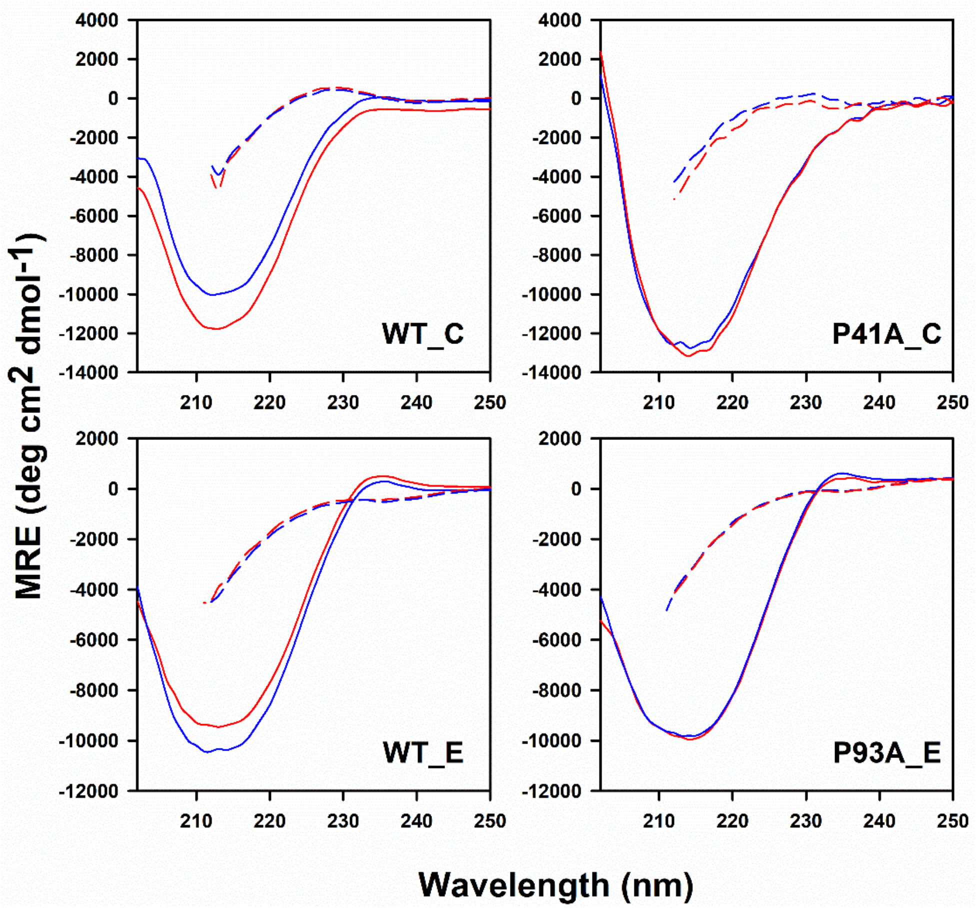


**Figure S1.** Far-UV CD spectra of the different mutant variants of MNEI. Spectra of the N (solid lines) and U (dashed lines) states are shown for the TNB-labeled (red lines) and unlabeled (blue lines) proteins. Panels WT_C and WT_E have been adapted and modified from reference 6.

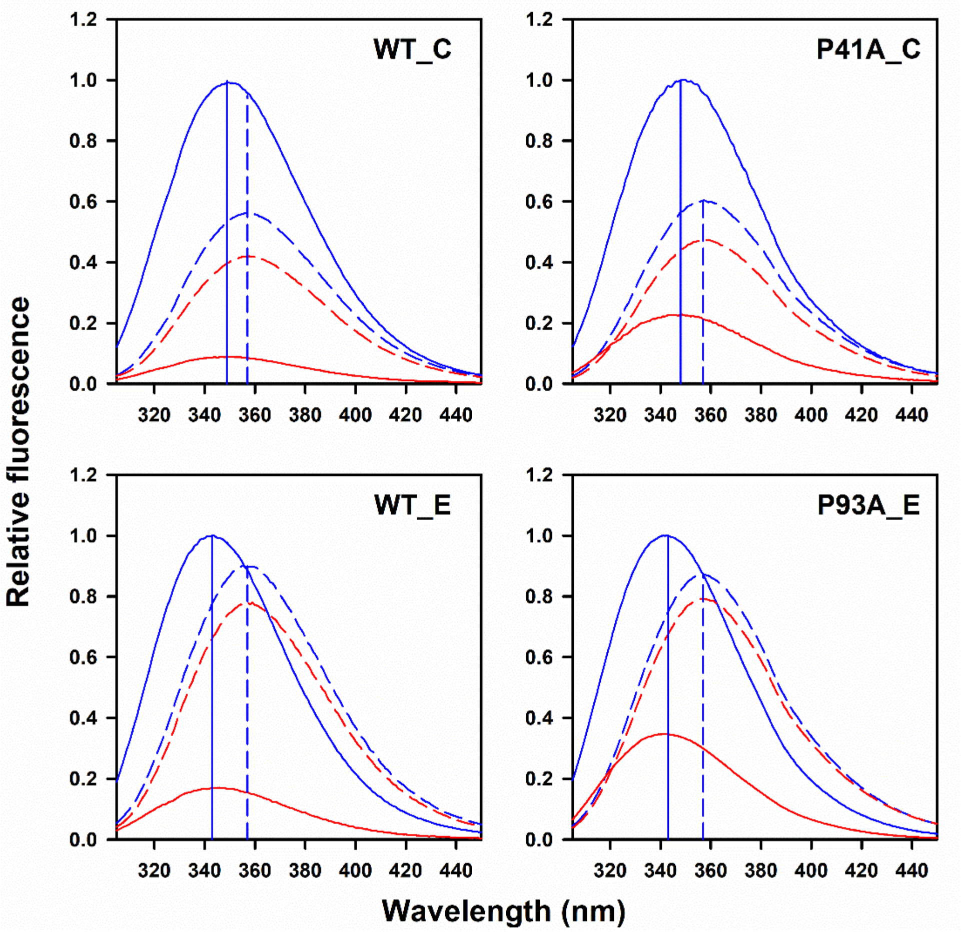


**Figure S2.** Fluorescence emission spectra of the different mutant variants of MNEI. The excitation wavelength was 295 nm. The solid and dashed curves represent the spectra of the native and unfolded states, respectively, for the TNB-labeled (red) and unlabeled (blue) proteins. The solid and dashed vertical lines indicate the peak positions of the spectra of the native and unfolded unlabeled proteins, respectively. Panels WT_C and WT_E have been adapted and modified from reference 6.

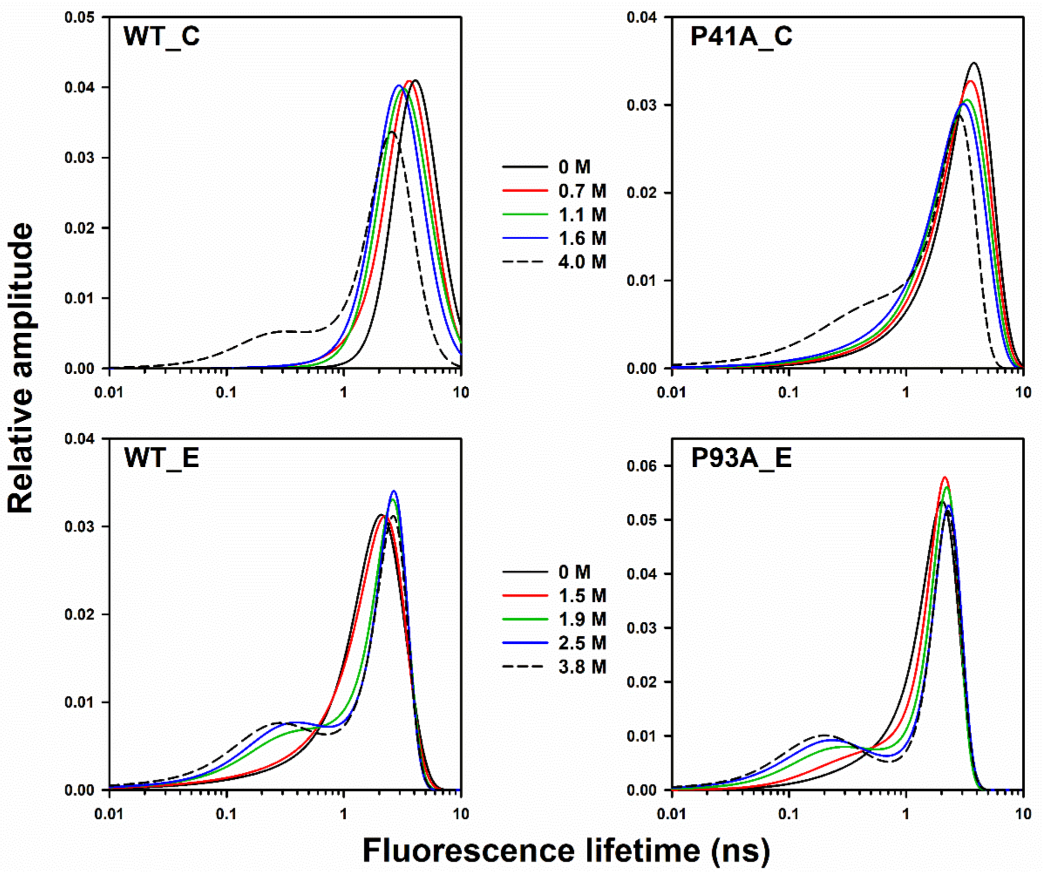


**Figure S3.** MEM-derived fluorescence lifetime distributions of the different mutant variants of MNEI at varying concentrations of GdnHCl. The colors of the lines represent the GdnHCl concentrations, as indicated. The data in panels WT_C and WT_E are from reference 6.

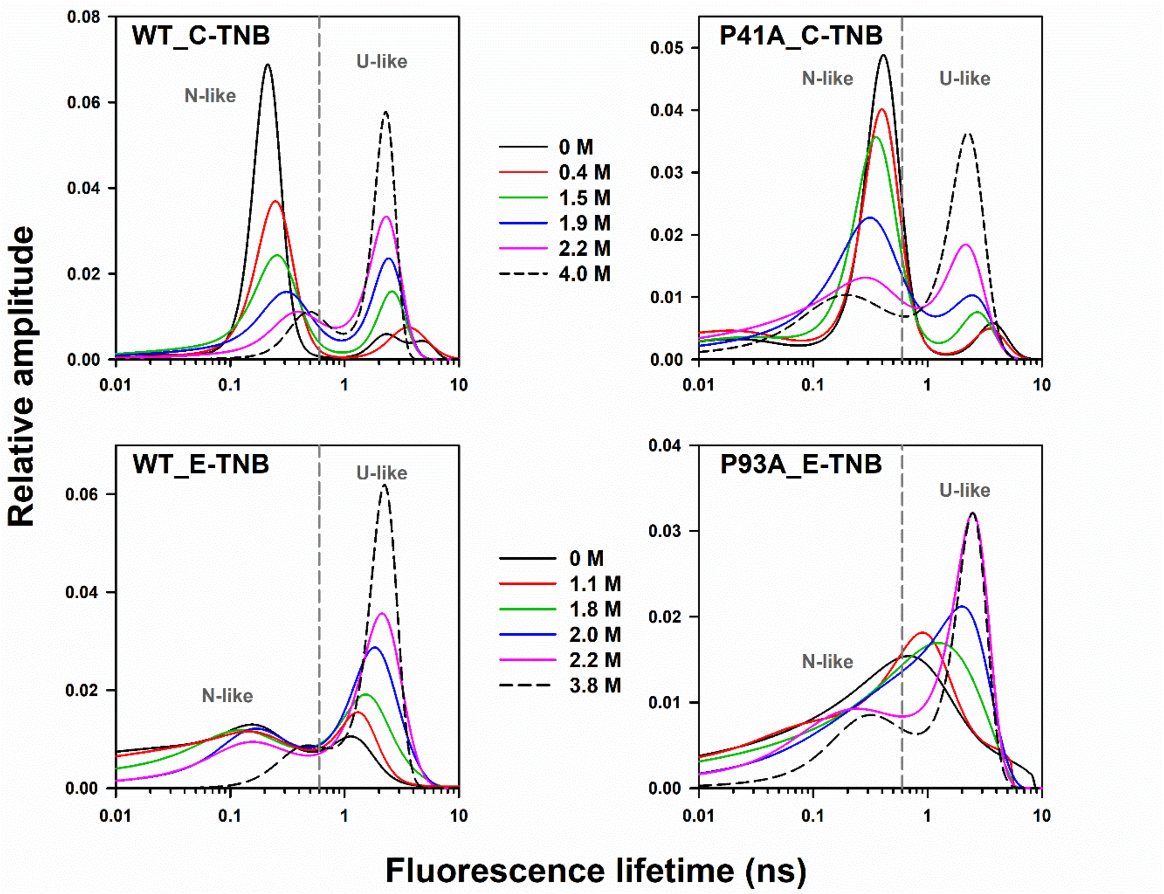


**Figure S4**. MEM-derived fluorescence lifetime distributions of different TNB-labeled variants of MNEI at varying concentrations of GdnHCl. The colors of the lines represent different GdnHCl concentrations, as indicated. The data in panels WT_C-TNB and WT_E-TNB are from reference 6.

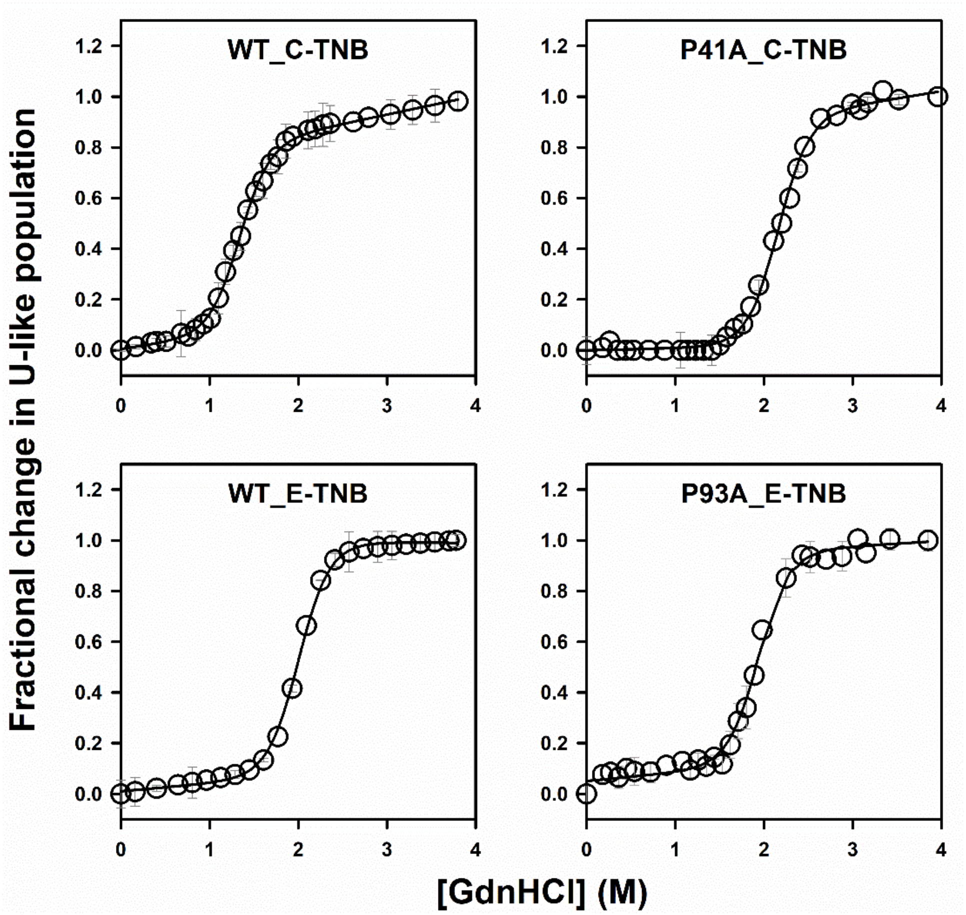


**Figure S5.** Fractional change in the U-like population calculated from the relative sum of amplitudes of the MEM distributions at different concentrations of GdnHCl. The fits to the two-state, N ↔ U unfolding model gave values for $\Delta G_U$ of 4.6 ± 0.3, 7.1 ± 0.2, 6.9 ± 0.1, and 6.7 ± 0.2 kcal $mol^{-1}$ for WT_C-TNB, P41A_C-TNB, WT_E-TNB and P93A_E-TNB, respectively. The mid-points ($C_m$) of the unfolding transitions for WT_C-TNB, P41A_C-TNB, WT_E-TNB and P93A_E-TNB are at 1.41 ± 0.08, 2.10 ± 0.04, 2.03 ± 0.02, and 1.97 ± 0.04 M GdnHCl, respectively. The error bars represent the spread in the data obtained from two independent experiments. Panels WT_C-TNB and WT_E-TNB have been adapted and modified from reference 6.

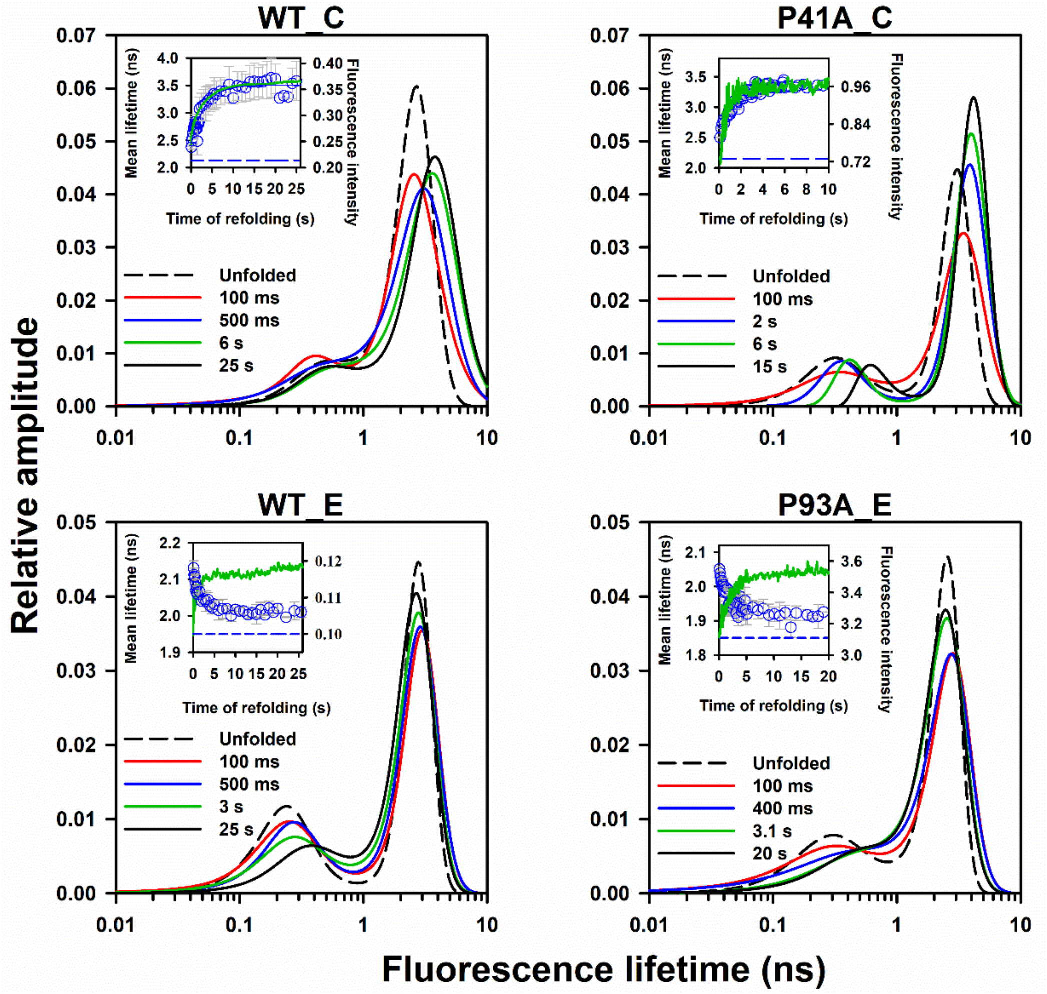


**Figure S6.** Kinetics of folding of the unlabeled MNEI variants in 0.4 M GdnHCl. The four panels correspond to different unlabeled variants (as indicated on the top of each panel). The dashed fluorescence lifetime distribution in each panel is of the U state. Distributions corresponding to different times of the folding reaction are shown in different colours as described for each panel. The inset in each panel represents the kinetics of folding monitored using simultaneous measurements of fluorescence intensity (green trace) and mean lifetime (blue circles) changes. The dashed blue line in the inset of each panel represents the mean lifetime of the U state. The error bars represent the standard errors of measurements from at least two independent double kinetics experiments. Panels WT_C and WT_E have been adapted and modified from reference 3.

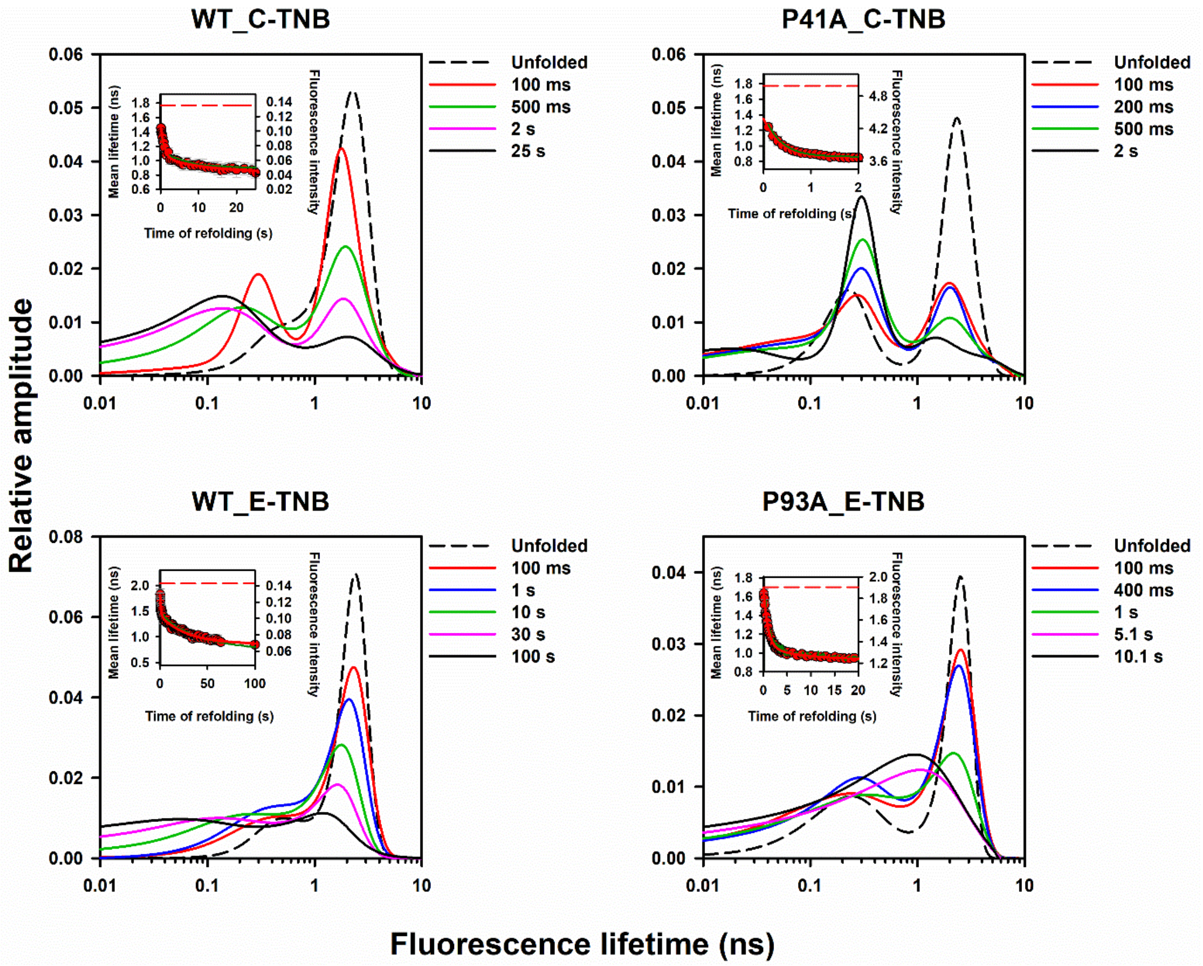


**Figure S7.** Kinetics of folding of the TNB-labeled mutant variants of MNEI in 0.4 M GdnHCl. The four panels correspond to different TNB-labeled mutant variants (as indicated on the top of each panel). The dashed fluorescence lifetime distribution in each panel is of the U state. Distributions corresponding to different times of the folding reaction are shown in different colours as described for each panel. The inset in each panel represents the kinetics of folding monitored using simultaneous measurements of fluorescence intensity (green trace) and mean lifetime (red circles) changes for different mutant variants of MNEI. The dashed red line in the inset of each panel represents the mean lifetime of the U state. The error bars represent the standard errors of measurements from at least two independent double kinetics experiments. Panels WT_C-TNB and WT_E-TNB have been adapted and modified from reference 3.

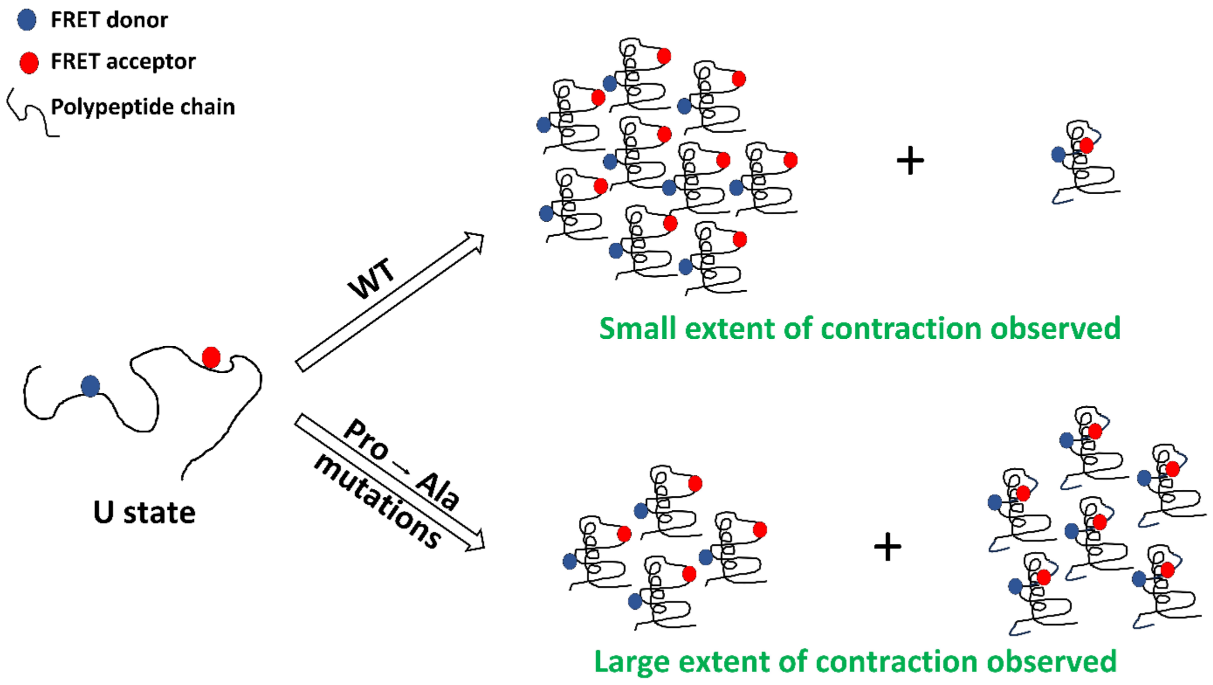


**Figure S8.** Schematic illustrating the effect of Pro to Ala mutations on the heterogeneity of an intermediate ensemble during the folding of MNEI. The mutations shift molecules toward more compact sub-populations, while the extent of structure formed remains largely unaffected. Symbols for donor, acceptor, and polypeptide chain are shown on the left. Numbers of molecules are illustrative and not exact.

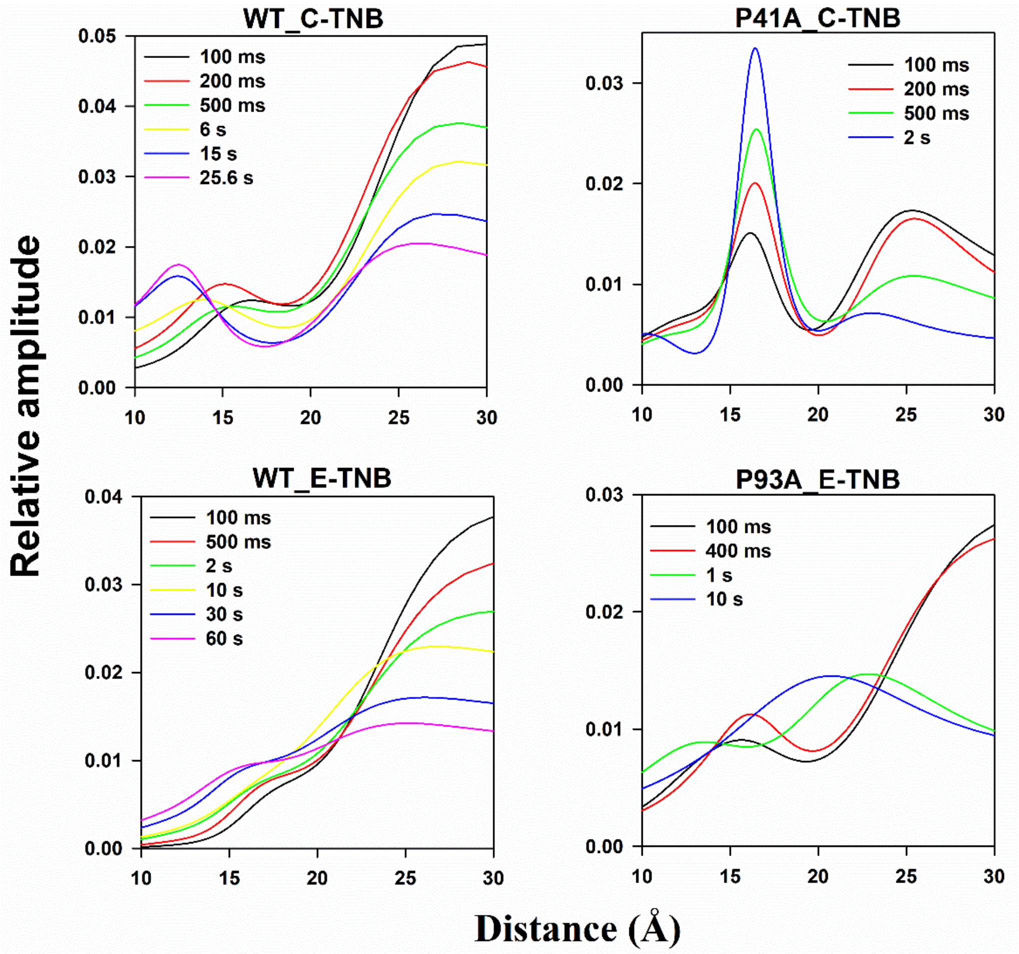


**Figure S9.** MEM-derived distance distributions at different times of folding of the different labeled variants of MNEI. In each case, the distributions have multiple cross-over points, indicative of a non-two state transition. Distributions corresponding to different times of folding are shown in different colours, as described for each panel. The y-axis in each panel represents the relative amplitude normalized to the sum of amplitudes for each distribution to make the total population fraction equal to 1. Panels WT_C-TNB and WT_E-TNB have been adapted and modified from reference 5.

**Table S1.** Thermodynamic parameters obtained from fluorescence-monitored equilibrium unfolding measurements for the different mutant variants of MNEI at pH 8 and 25°C.

| Protein variant | Free energy of unfolding, ($\Delta G_U$) kcal mol$^{-1}$ | Mid-point of unfolding,($C_m$) M |
|---|---|---|
| WT_C | 6.3 ± 0.1 | 1.86 ± 0.01 |
| WT_C-TNB | 4.9 ± 0.2 | 1.46 ± 0.06 |
| P41A_C | 5.4 ± 0.1 | 1.62 ± 0.02 |
| P41A_C-TNB | 6.9 ± 0.1 | 2.08 ± 0.07 |
| WT_E | 6.7 ± 0.4 | 1.97 ± 0.13 |
| WT_E-TNB | 6.9 ± 0.1 | 2.03 ± 0.01 |
| P93A_E | 6.6 ± 0.1 | 1.94 ± 0.04 |
| P93A_E-TNB | 6.8 ± 0.04 | 2.03 ± 0.01 |
